# Brain structure and function predict adherence to an exercise intervention in older adults

**DOI:** 10.1101/2021.07.30.454451

**Authors:** Timothy P. Morris, Agnieszka Burzynska, Michelle Voss, Jason Fanning, Elizabeth A. Salerno, Ruchika Prakash, Neha P. Gothe, Susan Whitfield-Gabrieli, Charles H. Hillman, Edward McAuley, Arthur F. Kramer

**Affiliations:** Department of Psychology, Northeastern University, Boston, MA, United States; Department of Human Development and Family Studies, Colorado State University, USA; Dept of Psychology, University of Iowa; Department of Health and Exercise Science, Wake Forest University, North Carolina, USA; Division of Public Health Sciences, Department of Surgery, Washington University School of Medicine in St. Louis, St. Louis, MO; Department of Psychology, Ohio State University; Beckman Institute for Advanced Science and Technology, University of Illinois at Urbana-Champaign, United States; Department of Kinesiology and Community Health, University of Illinois at Urbana-Champaign, Urbana, IL, United States; McGovern Institute for Brain Research, Department of Brain and Cognitive Sciences, Massachusetts Institute of Technology, Cambridge, MA, United States; Department of Physical Therapy, Movement, and Rehabilitation Sciences, Northeastern University, Boston, MA, United States

## Abstract

Individualized and precision medicine approaches to exercise for cognitive and brain health in aging have the potential to improve intervention efficacy. Predicting adherence to an exercise intervention in older adults prior to its commencement will allow for adaptive and optimized approaches that could save time (no need to demonstrate failure before changing course) and money (cost of de-implementing approaches that do not work for certain individuals) which ultimately could improve health outcomes (e.g., preventative medicine approaches prior to the onset of symptoms). Individual differences in brain structure and function in older adults are potential proxies of brain and brain reserve or maintenance and may provide strong predictions of adherence. We hypothesized that brain-based measures would predict adherence to a six-month randomized controlled trial of exercise in older adults, alone and in combination with psychosocial, cognitive and health measures. In 131 older adults (aged 65.79 ± 4.65 years, 63% female) we found, using regularized elastic net regression within a nested cross-validation framework, that brain structure (cortical thickness and cortical surface area) in somatosensory, inferior temporal, and inferior frontal regions and functional connectivity (degree count) in primary information processing (somatosensory, visual), executive control, default, and attentional networks, predicted exercise adherence (R^2^ = 0.15, *p* < 0.001). Traditional survey and clinical measures such as gait and walking self-efficacy, biological sex and perceived stress also predicted adherence (R^2^ = 0.06, *p* = 0.001) but a combined multimodal model achieved the highest predictive strength (R^2^ = 0.22, *p* <0.001). Neuroimaging features alone can predict adherence to a structured group-based exercise intervention in older adults which suggests there is substantial utility of these measures for future research into precision medicine approaches. The best performing model contained multimodal features suggesting that each modality provided independent relevant information in the prediction of exercise adherence.

## 1. Introduction

Physical activity and structured exercise have received a lot of attention as potential efficacious interventions to improve or maintain brain health with advancing age (Gomes-Osman et al., 2018; Kelly et al., 2014; Kramer et al., 2003). Despite millions of government and private dollars being spent on understanding how physical activity can improve or maintain brain function across the lifespan, over a third of the US population do not engage in sufficient physical activity (Du et al., 2019; Guthold et al., 2018; Kohl et al., 2012), a statistic which continues to increase (Du et al., 2019; Guthold et al., 2018). This physical inactivity pandemic (Andersen et al., 2016; Kohl et al., 2012; Pratt et al., 2020) is estimated to cost private and public health-care systems $53.8 billion per year and a further $13.7 billion in productivity losses due to physical inactivity-related deaths each year (Ding et al., 2016, 2017).

Previous research on understanding the adoption and maintenance of physical activity has leveraged psychological and psychosocial theories (Brand & Cheval, 2019). In experimental studies, several psychosocial, behavioral and demographic measures were shown to correlate with exercise adherence, such as self-efficacy (Garcia & King, 1991; McAuley et al., 2011), self-regulation (De Bruin et al., 2012; Dishman et al., 2014), social support, perceived benefits and biological sex (males adhered more than females) (Cadmus-Bertram et al., 2014), higher baseline physical activity outside of the intervention (Arkkukangas et al., 2018) as well as depression, fatigue and general perceived health (Flegal et al., 2007). Additionally, greater cognitive resources, particularly executive functions, known to depend on cortical structural integrity (Burzynska et al., 2011), have been associated with engagement in exercise (Cheval, Daou, et al., 2020; Cheval, Orsholits, et al., 2020). Based on these results, a multicomponent approach to physical activity behavior change has been proposed (Lachman et al., 2018). While theory-based programs are generally successful, optimizable, and adaptable precision medicine approaches will improve cost-efficiency of interventions and health outcomes for individuals who show poor adherence to one-size fits all approaches. That is, predicting from a set of variables, who is more or less likely to adhere to an intervention from the outset will allow the practitioner to provide alternative and individualized interventions prior to individuals having to either demonstrate poor adherence or present with deteriorating health.

Within the intervention setting, several recent studies have demonstrated that pre-intervention brain structure, known to be critical in supporting executive functions (Burzynska et al., 2011; Tadayon et al., 2020), is associated with individual differences in adherence to exercise interventions. Specifically, regions of the prefrontal, temporal and somatosensory cortices have been associated with adherence to each respective exercise intervention (Best et al., 2017; Gujral et al., 2018). Relatedly, several reviews and observational studies have suggested that the relationship between exercise and the brain is bi-directional (Audiffren & André, 2019; Cheval, Orsholits, et al., 2020). This hypothesis leverages the concepts of cognitive reserve and brain maintenance (Stern et al., 2020), such that there is a circular nature between greater cognitive and brain resources and higher participation in complex exercise behaviors, which in turn helps with the maintenance and upkeep of cognitive and brain health. While rarely measured directly, proxy measures of brain reserve and brain maintenance include brain structure and brain functional connectivity (for a review see: Stern et al., 2019). In terms of function, numerous applications of functional and anatomical connectivity have led to the observation that the brain is organized into large-scale functional networks (Yeo et al., 2011). A number of these networks, such as the frontoparietal control network (FPCN), the default mode network (DMN) and the dorsal attention network (DAN) change with age (Grady et al., 2016) and are thought to be particularly important in age-related cognitive decline (Marek & Dosenbach, 2018; Ng et al., 2016). These networks also subserve internally and externally directed cognition (Spreng et al., 2013), including broadly defined executive function. Given the heterogeneity of these brain networks and their importance in higher-order cognition, individual patterns of functional connectivity in older adults may be predictive of exercise adherence. Indeed, prior work on mindfulness and meditation have demonstrated that functional connectivity within the default mode network and in frontal and temporal nodes are strong predictors of adherence (Saghayi et al., 2020).

The term “prediction” is used in a number of ways in the literature and can commonly refer to either the correlation of one variable in a group at one point in time with another variable in that same group at another point in time (within-sample correlation) or it can refer to a generalizable model that makes predictions on out-of-sample participants (Gabrieli et al., 2015). Prior studies on psychosocial and behavioral “predictions” of exercise adherence have typically used correlation and therefore their results are likely overly optimistic (Yarkoni & Westfall, 2017). This overestimation and poor generalizability is compounded further when one considers they only explain relatively small amount of variance (<20%) in adherence (Rejeski et al., 2007; Rhodes et al., 2001). Consequently, calls have been made to introduce prediction statistics into psychological and cognitive neuroscience research to improve the generalizability of the results, specifically when using brain-based metrics (Gabrieli et al., 2015). In the case of exercise interventions for example, being able to successfully predict adherence to exercise interventions with generalizable results prior to engagement in the intervention would allow for optimization or individualization of an alternative intervention for those who are predicted to poorly adhere. For example, by providing those who are predicted to adhere poorly with a health coach (Olsen & Nesbitt, 2010) or just-in-time messaging (Nahum-Shani et al., 2018), paving the way towards efficacious precision medicine approaches in health-based settings, when time can rarely be lost, and cost-efficiency is of upmost importance. This is also important as engagement in effective exercise interventions may lead to successful maintenance of physical activity after cessation of the intervention (Madigan et al., 2021) and so predicting adherence to the exercise intervention may also provide insights into sustained exercise behaviors.

In a secondary analysis of data from a randomized control trial of exercise in older adults, the objectives of this study were to predict adherence to a 6-month exercise intervention and an active control in a structured and supervised group-based intervention. We hypothesized that multiple metrics of brain structure and function in frontoparietal control, default mode and dorsal attention networks and regions (prefrontal, temporal, parietal) would predict adherence to the exercise intervention and that these measures would augment traditional behavioral and psychosocial measures.

## 2. Methods

### 2.1 Participants

This study is a secondary analysis of data from participants who participated in a six-month randomized controlled exercise trial (clinical study identifier: NCT01472744, November 16, 2011). The study procedures were approved by the University of Illinois Institutional Review Board and written informed consent was obtained from all participants prior to any research activities. Healthy but low active older adults were recruited in Champaign County, Illinois. Two hundred and forty-seven (169 women) low-active older adults met inclusion criteria for the initial clinical trial. Of which, 165 underwent structural and functional magnetic resonance imaging (MRI) at baseline and 131 had complete data across all variables and were included in this analysis (see further exclusion criteria based on adherence below). Participants in the initial trial were randomized to one of four intervention conditions; a walking intervention, the same walking intervention plus a dietary supplement designed to enhance lean muscle mass, a dancing intervention and an active control consisting of a stretching and toning intervention. All intervention groups met for approximately one hour, three times per week for six months and for the purpose of this analysis we analyzed our main outcome (adherence) with all four conditions combined. A one-way analysis of variance revealed no significant differences in adherence between the conditions (F_1,129_ =1.11, *p* = 0.294) and no significant differences in baseline characteristics between conditions existed (supplementary table 1). Further we replicate our results in the most common intervention conditions (walking) only as a sensitivity analysis with a reduced sample size (n=59) to ensure our results were not affected by condition assignment (supplementary material). Initial inclusion criteria included being between the ages of 60 and 80 years old, free from psychiatric and neurological illness including no history of stroke, transient ischemic attack, or head trauma, scored > 23 on the Mini-Mental State Exam, > 21 on a Telephone Interview of Cognitive Status questionnaire, < 10 on the Geriatric Depression Scale, at least 75% right-handed based on the Edinburgh Handedness Questionnaire (a criterion related to functional magnetic resonance imaging (MRI) analyses), demonstrated normal or corrected-to-normal vision of at least 20/40 and no color blindness, screened for safe participation in an MRI environment (e.g., no metallic implants that could interfere with the magnetic field or cause injury and no claustrophobia) and reported to have participated in no more than two bouts of moderate exercise per week within the past six months (with the goal of recruiting low active older adults). Table 1 contains complete characterization of the study participants included in this analysis. All methods were carried out in accordance with the Declaration of Helsinki.

**Table 1.** Participant Characteristics.

|  |  |
| --- | --- |
| N | 131 |
| Age (mean (SD)) | 65.79 (4.65) |
| Gender: Female (%) | 63 (71.6) |
| Race (%) |  |
| White | 79 (89.7) |
| African American or black | 7 (8) |
| Asian | 2 (2.3) |
| Years of education (mean (SD)) | 15.89 (2.66) |
| Adherence (%) | 81 (12) |

### 2.2 Adherence

Our primary outcome, adherence, was modelled as percentage attendance to the weekly intervention sessions. These supervised sessions were scheduled three times per week and lasted approximately for one hour. The number and frequency of the sessions were consistent across all four conditions. For this analysis, participants (N=34) who dropped-out of the study (i.e., did not complete the intervention) were excluded, as were those who failed to attend at least half of the intervention sessions (n= 12) as our aim was to capture and predict variations in adherence to the entire 6-month intervention.

### 2.3 Psychosocial, physical function and activity, health, and cognitive features

A comprehensive battery of psychosocial, physical and cognitive assessments was completed by each participant pre-intervention. A complete table of these assessments is found in the supplementary material. Eighty-four assessments were initially included in this analysis. In brief, these assessments included self-reported psychosocial questionnaires gauging participants self-efficacy (global, exercise and gait self-efficacy), leisure time activity, perceived sleep quality, anxiety, depression and self-worth/esteem, barriers to exercise, self-regulation, stress, loneliness and subjective memory. All psychosocial measures were taken at week one or in some instances repeated at week three. A set of physical function tests were also collected at week one, which included a stair climb test, arm curl, sit and reach and back scratch. Seven days of accelerometry capturing objective measures of time spent sedentary, time spent in light or moderate-to-vigorous aerobic physical activity and average daily step counts were also collected. The procedures to capture, preprocess and validate these measures can be found in previous publications (Burzynska et al., 2020; Fanning et al., 2017). A measure of cardiorespiratory fitness from a complete cardiopulmonary exercise test was also included as were measures of body composition (see our previous works (Burzynska et al., 2014; Voss et al., 2016) for a detailed description of the methodology of these measures). Finally, a battery of neuropsychological tasks were completed in week one which included numerous assessments of vocabulary, abstract, inductive and visuospatial reasoning, memory and perceptual speed, taken from the Virginia Cognitive Aging Project (Salthouse & Ferrer-Caja, 2003). Intervention condition assignment was also included as a feature.

### 2.4 Neuroimaging features

#### 2.4.1 Magnetic resonance imaging: acquisition

Participants undertook an MRI scanning session in a three Tesla Siemens Trio Tim system with a 12-channel head coil. High-resolution structural MRI scans were acquired using 3D MPRAGE T1-weighted sequences (TR = 1900 ms; TE = 2.32 ms; TI: 900 ms; flip angle = 9°; matrix = 256 × 256; FOV = 230 mm; 192 slices; resolution = 0.9 × 0.9 × 0.9 mm; GRAPPA acceleration factor 2). T2*-weighted resting state echoplanar imaging (EPI) data was obtained with the following parameters: (6min,TR=2s,TE=25ms,flipangle=80°, 3.4 × 3.4 mm^2^ in-plane resolution, 35 4 mm-thick slices acquired in ascending order, Grappa acceleration factor = 2, 64 × 64 matrix). Structural and resting state functional images were acquired with these scanning parameters and the preprocessing and analyses of each respective modality are outlined in the following two sections.

#### 2.4.2 Structural MRI preprocessing and analyses

Cortical reconstruction and image segmentation and estimation of the cortical surface models was performed using the freely available FreeSurfer software v.5.3 (http://surfer-nmr.mgh.harvard.edu/). For preprocessing of the cortex a 3-dimensional surface model was created using the “recon-all” surface-based stream. Automated Talairach transformation and intensity normalization were followed by nonbrain tissue removal, tessellation of the gray and white matter boundary and automated topology correction. Finally, surface deformation enabled the detection of tissue boundaries; grey–white and grey–CSF borders. The cortical surfaces were then inflated and registered to a spherical atlas that used individual cortical folding patterns to match cortical geometry across participants. *Cortical thickness* was calculated at each vertex in the cortex as a measure of the distance between the white and pial surfaces and *cortical surface area* was calculated by averaging the area of all faces that meet at a given vertex on the white matter surface. We chose to analyze cortical thickness and surface area separately given their genetic independence and sensitivity to clinical and aging outcomes (Storsve et al., 2014). Automatic labelling per the Desikan-Killany cortical parcellation scheme was performed and average cortical thickness and cortical surface area were calculated within each parcellation resulting in 136 structural features (68 features per modality) to be used for feature selection.

#### 2.4.3 Resting state functional connectivity preprocessing and analyses

Preprocessing of the functional resting state data was performed using the CONN-toolbox v.19c (Whitfield-Gabrieli & Nieto-Castanon, 2012), relying upon SPM v.12 (Wellcome Department of Imaging Neuroscience, UCL, London, UK) in MATLAB R2019a (The MathWorks Inc, Natick, MA, USA). The default preprocessing pipeline implemented in CONN was performed which consists of the following steps: functional realignment and unwarping, slice timing correction, outlier identification, segmentation (into grey matter, white matter and cerebrospinal fluid tissue) and normalization into standard Montreal Neurologic Institute (MNI) space with 2mm isotropic voxels for functional data and 1mm for anatomical data, using 4^th^ order spline interpolation. Finally, functional scans were spatially smoothed using a 6mm Gaussian kernel. During the outlier detection step, acquisitions with framewise displacement above 0.9mm (per several prior publications in studies with older adults who are more susceptible to movement within the scanner (Fitzhugh et al., 2019; Pistono et al., 2021; Vieira et al., 2020) or global BOLD signal changes above five standard deviations were flagged as potential outliers using the Artefact Detection Tools (www.nitrc.org/projects/artifact_detect). Two participants were removed from the final analyses for having >40 volumes flagged. This cut off was determined based on preserving at least five minutes of scanning time (Van Dijk et al., 2009). Additionally, mean framewise displacement was calculated via the Jenkinson method (Jenkinson et al., 2002) and regressed out of the final analysis (see statistical analysis section below). This was done to be over-conservative given previous studies have shown high degree of motion-behavior correlations (Siegel et al., 2017), despite the fact that no motion variable was significantly correlated with adherence outcomes in our study (all p > 0.1). Denoising of the functional data was performed using a component-based correction method, CompCor (Behzadi et al., 2007) and temporal band-pass filtering (0.01-0.1Hz) to remove physiological, subject-motion and outlier artefacts. Linear regression was used to remove the effects of these artifacts on the BOLD time series for each voxel and each subject taking into account noise components from voxels within white matter and cerebrospinalfluid, estimated subject-motion parameters (three rotation and three translation parameters and six other parameters representing their first order time derivatives), scrubbing and constant and first-order linear session effects.

To prepare the functional data for feature selection, we parcellated the functional scans into a medium resolution 300 region (node) atlas with a 17-network parcellation scheme (Schaefer et al., 2018). This choice was taken to strike a balance between biological resolution and dimensionality. Mean BOLD activity was calculated within each node and transformed into a 300 by 300 correlation matrix where the timeseries at each node was correlated with that of every other node. This matrix then underwent Fisher’s Z transformation. Functional brain connectivity was summarized using degree count, a graph theory metric which represents whole brain connection density for each node, per a previous publication (Vogel et al., 2018). This metric reduces the dimensions of the functional connectivity data into a sparse matrix that better represents real world graphs (Power et al., 2013). Degree count for each participant was calculated at several thresholds using a fixed network cost (keeping the strongest 15%, 20% and 25% of connections) and a final threshold was chosen through cross-validation (see statistical analysis section). Degree count at each node here represents a single measure of the sum of connections between that node and every other node in the cortex characterizing its degree of connectedness within the cortical gray matter.

### 2.5 Power analysis

We performed a power analysis (in R using the “pwr” package) on our sample size to ensure sufficient power was gained to detect a true effect. Based on our sample size (N = 131) and an assumed type I error rate of 0.05, we calculated an estimated 96% power to detect an effect size of 0.22 from a general linear model with 38 covariates (multimodal model). In the reduced sample size of N=59 for the sensitivity analysis (supplementary material) this power fell to 60% for the same effect size.

### 2.6 Statistical analysis

To predict adherence to the exercise interventions we used Elastic net regression, a regularized (penalized) regression method within a nested cross-validation procedure. Elastic net aims to avoid overfitting by producing less-complex models via applying both a L1 and L2 norm penalty to calculate the coefficients of each feature. Elastic net is a linear combination of both Ridge regression (L2 norm) and least absolute selection and shrinkage operate (Lasso) regression and as such strikes a balance between shrinking coefficients towards zero (Ridge) and in the case of features that have no predictive value, to exactly zero (Lasso), relative to the maximum likelihood estimates. The predictors with non-zero coefficients are therefore interpreted as those that contained predictive information and contributed to the final model of predicting our outcome. This model is particularly useful in cases where correlated features are present (i.e., neuroimaging features and some psychosocial measures) as the combination of both the L1 and L2 penalties will maintain groups of correlated features in the model (whereas the L1 penalty (Lasso) alone would remove all but one of the correlated features). The amount of shrinkage is determined via tuning of two hyperparameters λ_1_ and λ_2_. Results of the optimal hyperparameters for each model are found in Table 2. In this analysis we used a nested cross-validation procedure where the dataset is split into a 10-fold outer loop (to evaluate model performance) and a 10-fold inner loop (to tune the hyperparameters using grid search within each inner fold). Nested-cross validation avoids optimization bias that simple cross-validation could potentially suffer from when using the same folds to both tune the hyperparameters and test the prediction performance on. In this case, the folds were kept consistent across each model by setting the same random seed and all variables were centered and scaled within each inner loop and applied to the outer folds. First, to select the optimal degree count matrix threshold, models were trained over each threshold density and the model that predicted the left-out outer folds with the smallest root mean squared error (RMSE) was used in the final models. In the final models, a manual feature preselection was performed by removing features that didn’t correlate with adherence at p<0.1 (p<0.1 was selected to maintain only features that contained potentially predictive information) and age, biological sex and mean framewise displacement (functional connectivity only) were regressed out of the features. Three separate models were trained: (1) imaging features only, (2) Psychosocial, physical function and activity, health, and cognitive features only and (3) multimodal features containing all features. In the case of model 2, the combination of nested cross-validation and Elastic net failed to predict adherence in the left-out outer folds, potentially due to the small number of features in the model (p=6) and as such, for the non-imaging model only we ran a simple 10-fold cross-validation model using Ridge regression (L1 norm penalty only). Model performance (prediction on the left-out folds) was calculated as the squared correlation between the predicted and observed values (R^2^). Other performance metrics included RMSE, which measures the average prediction error as the average difference between the observed and predicted values and the mean absolute error (MAE) as the average absolute difference between the observed and predicted values. RMSE and MAE are related with MAE being less sensitive to outliers and the lower the value the better the model performance. For R^2^, a higher value equates to better performance. To assess the significance of the R^2^ prediction performance, 1000 non-parametric permutations were performed, and the p-value of the permutation tests was calculated as the proportion of sampled permutations that are greater or equal to the true prediction correlation. All statistics were performed in RStudio version 3.6.3 using “dplyr”, “purr”, “penalized” and “pensim” packages. Figures were generated using “ggplot2”.

**Table 2.** Model performance metrics.

| Model | $R^2$ | RMSE | MAE | P-value |
| --- | --- | --- | --- | --- |
| Multimodal | 0.22 | 0.10 | 0.09 | <0.001 |
| Imaging | 0.15 | 0.11 | 0.09 | <0.001 |
| Psych/cog/health | 0.06 | 0.12 | 0.10 | 0.001 |

Performance metrics derived from nested cross-validation where the optimal hyperparameters for Elastic net regression were tuned in an inner loop and used to predict adherence in the left out outer loop. The p-value for each model is derived by comparing the actual $R^2$ against a null distribution derived from 1000 non-parametric permutations.

## 3. Results

Table 1 presents demographic details of the participants included in this analysis. Our sample consisted of mostly female, white, and educated participants and the mean adherence was 81%.

Table 2 contains prediction metrics for each modality using Elastic net models and Ridge regression for the cognitive/psychosocial only model. All models resulted in a significant prediction of adherence to the exercise interventions with the multimodal model performing the best, followed by the imaging and then the psychosocial, physical function and activity, health, and cognitive model (Table 2 and Figure 1).

**Figure 1.**
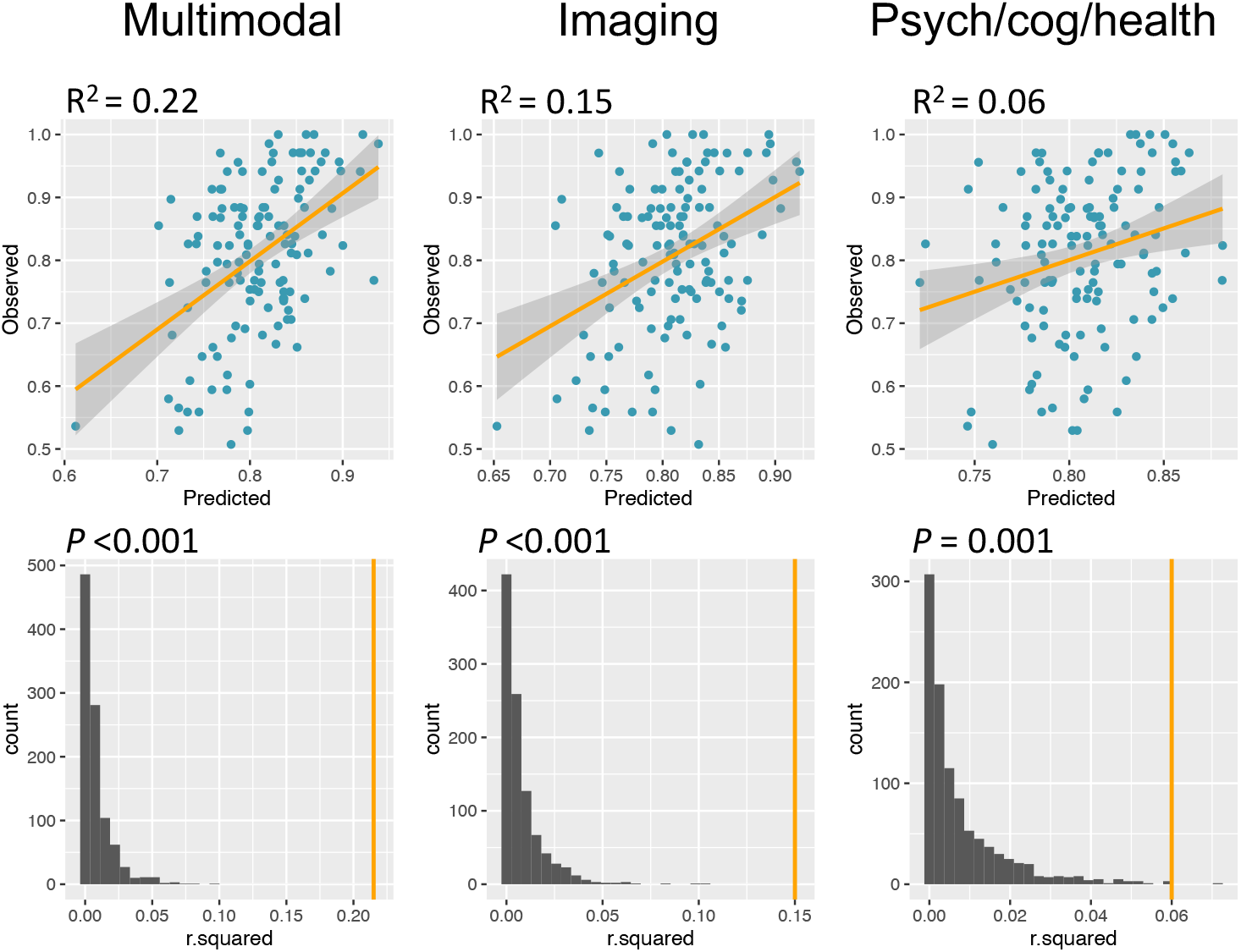
Prediction performance (R^2^) of each model (top row) and the significance test (*p*-value) of each model via 1000 permutations (bottom row). Top row: R^2^ represents the squared correlation between the predicted and observed adherence values via Elastic net regression with nested cross-validation. Bottom row: Null distribution of the prediction performance via 1000 non-parametric permutations. The orange vertical line represents the actual model performance (R^2^) and the *p*-value from the permutation tests is calculated as the number of sampled permutations that are greater or equal to the true prediction correlation.

Univariate correlations between those features selected after manual feature preselection and adherence is found in Table 3. Standardized coefficients for each feature selected in each model are found in Table 3. A summary figure of the multimodal model is found in Figure 2. For the functional connectivity features, the result broadly shows multiple distributed negative nodes in the somatomotor, visual, temporoparietal and ventral attention networks and multiple positive nodes distributed across DMN, DAN and FPCN predictive of adherence. For the structural features, distributed features across both hemispheres which were almost all positive for cortical thickness and negative for surface area were found to predict adherence. For the Psych/cog/health features, several psychosocial questionnaires (i.e., gait and walking self-efficacy, stress and sleep quality), biological sex and one neuropsychological test (i.e., letter comparison) were selected as predictive features.

**Table 3.**
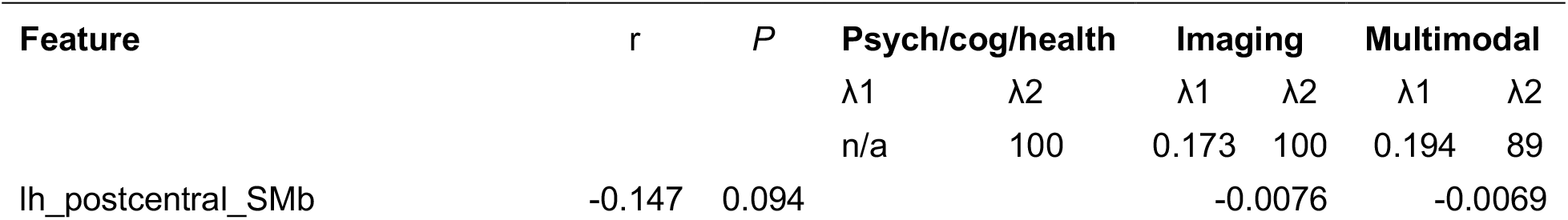
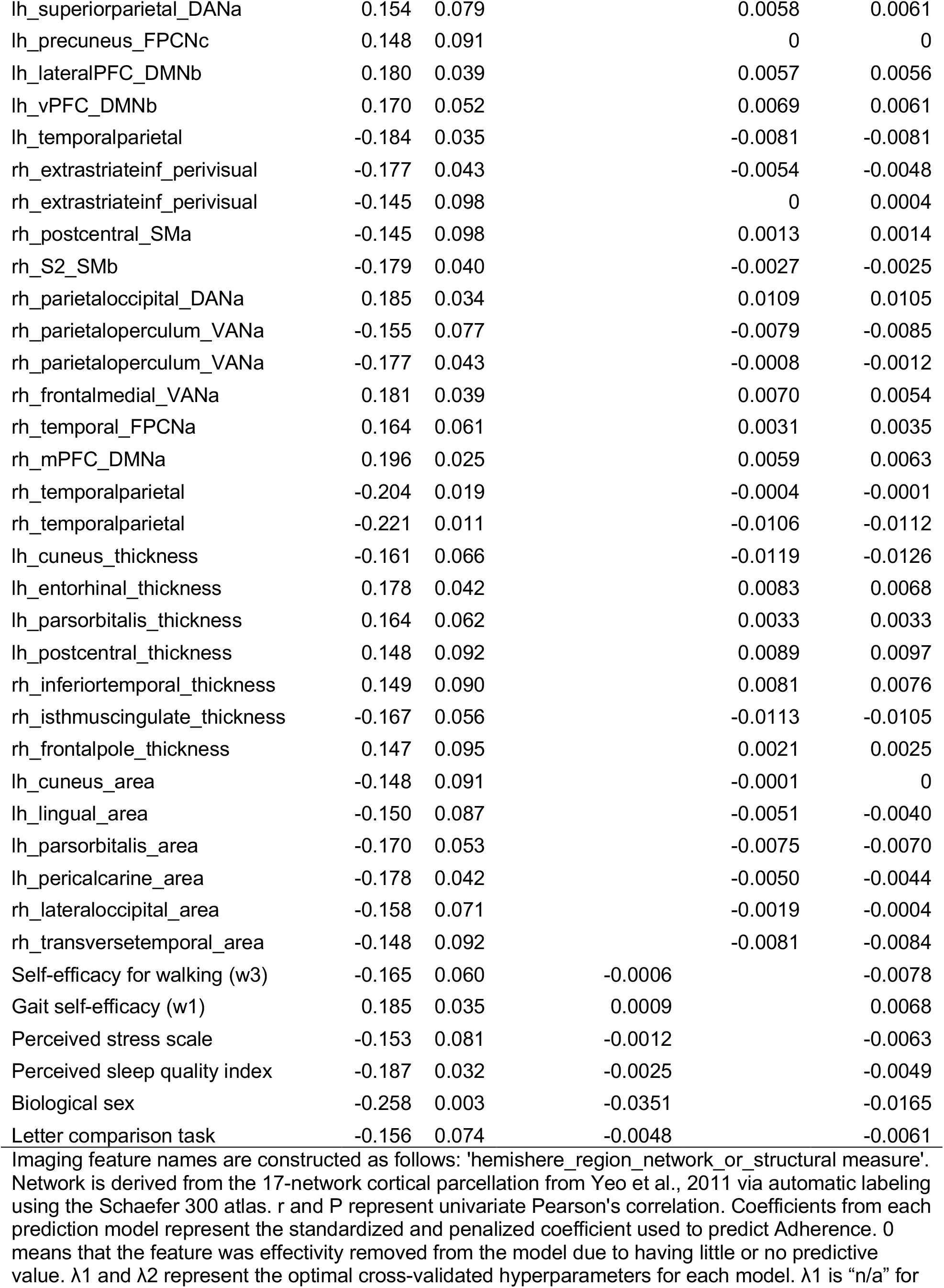
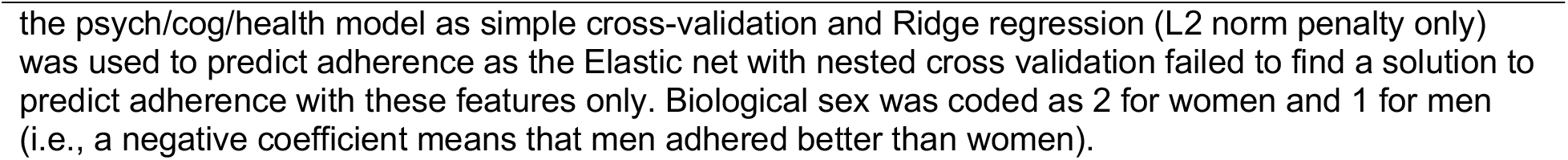
Univariate correlations and Elastic net feature selection

**Figure 2.**
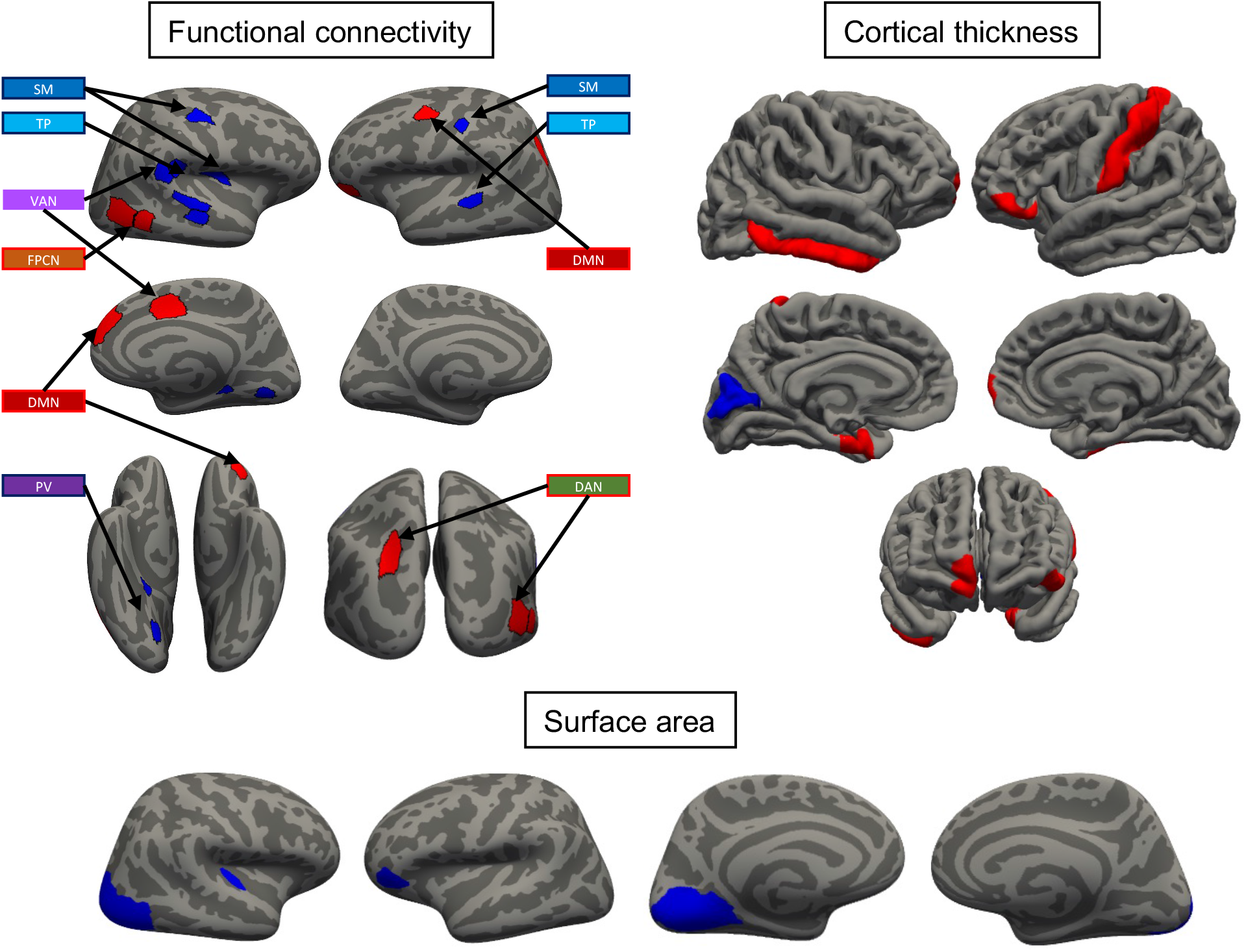
Summary figure depicting the selected features of brain structure and function in the multimodal model (best performing model). For the functional connectivity features (top left), each feature represents degree count (number of connections that each node has with every other node in the cortical parcellation). That is, red features (positive nodes) are interpreted as the higher the connectivity between that node and every other node in the cortex, the higher the adherence. Blue features (negative nodes) suggest that less connections between these nodes and the rest of the brain the higher the adherence. Mapping these regions to intrinsic resting state functional networks (17 Yeo networks via the Schaefer 300 atlas) reveals a broad pattern of positive nodes in the DMN, DAN and FPCN and negative nodes in primary information processing networks. In general, greater cortical thickness and lower cortical surface area in several disperse brain regions were predictive of adherence.

## 4. Discussion

In this study we demonstrated that brain-based measures of functional connectivity, cortical thickness and surface area predicted future adherence to a structured group-based exercise intervention in older adults. Using a machine learning framework that applies penalized regression with cross-validation we also replicated earlier studies indicating that aspects of self-efficacy and biological sex are predictive of adherence to exercise and that together, multimodal features provide the strongest predictive value.

Currently, our model performance (R^2^ of .22) remains far from having utility in a clinical setting. While we replicated the finding that traditional psychosocial and demographic measures predict adherence to exercise, the ability of neuroimaging features alone to predict adherence to a structured group-based exercise intervention suggests substantial utility of these measures for future research into precision medicine and adaptive intervention approaches. Additionally, the increase in model performance when combined suggests that multimodal features provide independent relevant information in the prediction of exercise adherence.

In a research setting, the application of prediction modelling could aid in the development of adaptive intervention strategies. For example, in those who can successfully engage in a theory-based intervention, positive results have been shown. However, successful engagement in long-term interventions is non-trivial and not all individuals will adhere. If one could predict these individuals from the outset, re-deployment of resources to tailor the intervention in these individuals with the goal of improving adherence and subsequent efficacy could be done. For example, our results could be leveraged in future prospective studies to test whether adaptive interventions based on pre-intervention characteristic and brain signatures (our predictions) result in better adherence and intervention efficacy. Additionally, successful engagement in an exercise intervention may also be related to long-term maintenance of exercise behaviors beyond the intervention (Madigan et al., 2021). Consequently, applying this type of prediction modeling to recent large population and cohort studies (e.g., UK Biobank or Human Connectome Project) with imaging and self-report physical activity data could help one determine the proportion and characteristics of individuals who would require additional tailoring when designing future exercise interventions.

Regarding our imaging analysis, we took a whole-brain approach to the prediction of exercise. Our results however could be leveraged to validate these results in future hypothesis-driven studies. For example, nodes with high global functional connectivity predictive of adherence in our study were found in the DMN, DAN and FPCN, three related networks that contribute to internally and externally directed cognition (Spreng et al., 2013) and in the implementation of executive control processes to maintain goals and inhibit distractions, respectively (Spreng et al., 2010; Vincent et al., 2008). Similarly, cortical structure in postcentral, ventrolateral/orbitofrontal, and inferior temporal regions have been shown to be associated with executive functions, especially in healthy aging (Burzynska et al., 2011). As such, direct study of the interplay between these networks at rest or during executive function tasks that engage these networks may provide stronger predictive utility.

Numerous randomized controlled trials of exercise in aging have demonstrated that exercise can improve (with mixed effect sizes) cognitive function and positively effect brain structure and functional connectivity (Erickson et al., 2019; Gomes-Osman et al., 2018; Northey et al., 2018). Yet the hypothesized bi-directional that the relationship between exercise and the brain and cognition is starting to be tested (Cheval, Orsholits, et al., 2020). In our current study, individual differences in proxy measures of cognitive and brain reserve (functional connectivity nodes, cortical thickness and surface area, respectively) within primary information processing networks and regions were predictive of exercise adherence. Prior exercise interventions have shown intervention-mediated increases in the functional connectivity of somatosensory networks in older adults (McGregor et al., 2018). Our results therefore potentially provide supportive evidence in favor of the hypothesis that the relationship between exercise and the brain is bi-directional (Loprinzi et al., 2013; Ponce & Loprinzi, 2018). Notwithstanding, our prediction models need to be tested in prospective and experimental studies to conclude a causal association.

Several limitations to this study mean that our results should be interpreted with reasonable caution. Firstly, the complexity of adherence to a 6-month exercise intervention may not be completely captured in a single variable representing the percentage attendance to the intervention settings and so additional explained variance not captured by our measures may be due to other reasons such as scheduling conflicts related to family, work or breaks due to discomfort. Second, it is of note that inferential guarantees regarding variable estimates (coefficients) in penalized regression models cannot be made. I.e., traditional p-values or confidence intervals for each estimate do not exist (Lockhart et al., 2014). Therefore, estimates produced by the elastic net model are biased and should not be interpreted as the population parameter. While advances in post-selection inference methods for the Lasso have been made ((Lee et al., 2016; Lockhart et al., 2014) which allow for more valid confidence intervals and significance testing for Lasso estimates), such methods do not exist for Elastic Net, the use which was important in our case given the collinearity of the predictors in our study. Third, our sample size was relatively small for machine learning. As such, testing on completely left-out subjects (test-train splits) was not feasible and so we attempted to perform the most rigorous and generalizable approach possible via nested cross-validation. Fourth, given the difficulty and costs associated with this type of research and the vast number of measures that were available in this unique dataset, replication of this prediction in a completely independent validation dataset is unavailable at this moment. Fifth, our participants were homogenous. Several factors have recently been highlighted as a reason why global physical inactivity rates continue to be low (Pratt et al., 2020) with one being that most of the research on understanding physical activity behaviors has occurred in high-income countries and so more work in diverse populations and low-income countries is needed to fully generalize this type of prediction. Lastly, our battery of neuropsychological tasks included executive function tests that measured constructs such as abstract, inductive and visuospatial reasoning rather than other executive tasks like inhibitory control that may be more related to exercise adherence (Cheval, Daou, et al., 2020).

Our results showed that the combination of psychosocial, cognitive and demographic, and multimodal imaging metrics can predict adherence to an exercise intervention in older adults and provide independent relevant predictive value. Prospective testing of these predictions and their validation will allow researchers and eventually clinicians to leverage them through personalized medicine approaches.

## Acknowledgments

We would like to thank, Anya Knecht, Susan Houseworth, Nancy Dodge, Hilly Tracy, Robert Weisshappel and all of the Lifelong Brain and Cognition and Exercise Psychology Laboratory graduate students and staff for their help in participant recruitment and data collection.

## Author contributions

TPM- Conceptualization, design, analysis, interpretation of data, manuscript writing, AB–data acquisition, substantial revision, MV-data acquisition, substantial revision, JF-data acquisition, substantial revision, ES - data acquisition, substantial revision, RP-data acquisition, substantial revision, NG-data acquisition, substantial revision, SWG-analysis, interpretation of data, CH – design, substantial revision, EM-Conceptualization, study design, substantial revision, AF-Conceptualization, study design, analysis, interpretation of data, substantial revision

## Ethics approval

The University of Illinois Institutional Review Board approved all procedures used in the study.

## Consent to participate

All participants gave written informed consent before participation in any study procedures, all of which conformed to the Declaration of Helsinki for research involving human subjects.

## Consent for publication and author responsibilities

All authors agree to the contents of this manuscript and give consent for its publication.

## Availability of data and materials

All data will be provided upon reasonable request to the corresponding author, without reservation.

