## Supplementary material for "Brain structure and function predict adherence to an exercise intervention in older adults"

**Brain-based measures improve the prediction of future adherence to exercise interventions in older adults.**

1. Optimal degree count threshold

To define the optimal degree count threshold to use in the final models in the main paper we ran separate elastic net models at each fixed network cost level (i.e., the strongest 15%, 20% and 25% of connections) within a similar nested cross validation framework whereby hyperparameters were tuned within 100 internal bootstrap samples and tested on 3 left-out outer folds. The average of the RMSE over the 3 outer folds was used to determine the optimal threshold to use (table 1).

| **Degree count fixed network threshold** | | | | |
|---|---|---|---|---|
| **15** | Fold1 | Fold2 | Fold3 | Average |
| | 0.1115787 | 0.1170570 | 0.1155732 | 0.1147363 |
| **20** | | | | |
| | Fold1 | Fold2 | Fold3 | |
| | 0.1143274 | 0.1202450 | 0.1098508 | 0.11480773 |
| **25** | | | | |
| | Fold1 | Fold2 | Fold3 | |
| | 0.09167418 | 0.12896172 | 0.12279353 | **0.11447648** |

2. Complete list of behavioral variables

| **Demographics** |
|---|
| Age |
| Biological sex |
| Education (years) |
| Employment status |
| Race |

| |
|---|
| Annual household Income |
| Intervention condition |
| **Cognition** |
| Digit symbol |
| Pattern comparison |
| Letter comparison |
| Word recall |
| Logical memory |
| Paired associates |
| Shipley abstract reasoning |
| Foamboards |
| Letter sets |
| Matrix reasoning |
| Paper folding test |
| Spatial relations |
| Vocabulary test |
| Picture vocabulary |
| Synonyms |
| Antonyms |
| **Psychosocial and lifestyle questionnaires** |
| Godin Leisure Time Physical activity questionnaire |
| Physical activity scale for the elderly |
| Satisfaction with life scale |
| Hospital anxiety and depression scale |
| Self-efficacy for walking scale (week 0 and week 3) |
| Lifestyle self-efficacy scale (week 0 and week 3) |

| |
|---|
| Gait self-efficacy scale (week 0 and week 3) |
| Sitting time questionnaire |
| Function self-efficacy scale (week 0 and week 3) |
| Barriers self-efficacy scale (week 0 and week 3) |
| Exercise self-efficacy scale (week0 and week 3) |
| Self-regulation scale |
| UCLA Loneliness Scale |
| Perceived Stress Scale |
| Physical activity enjoyment scale |
| Multidimensional outcome expectations scale |
| Life Orientations Scale |
| Big Five Personality Inventory |
| SF-12: Mental and Physical Health Status – |
| Frequency of forgetting |
| Memory Controllability Inventory |
| Rosenberg self-esteem questionnaire |
| Perceived sleep quality index |
| Social provisions scale |
| Social network inventory |
| Physical self-presentation scale |
| **Physical function and accelerometry** |
| Cardiorespiratory fitness ($V0_2$ max) |
| BMI |
| Resting heart rate |
| Average time spent sedentary (mins) |
| Average time in light intensity physical activity |

| |
|---|
| Average time in moderate to vigorous physical activity |
| Average step counts |
| Start climb test |
| Chair stand test |
| Arm curl test |
| Chair sit and reach test |
| Back scratch test |
| 8ft up and go test |
| Left and right leg stand test |

3. Demographics stratified by group

Supplementary table 3. Demographics stratified by group assignment

| | **Dance** | **Stretch/tone** | **Walk** | **Walk +** | ***p*** |
|---|---|---|---|---|---|
| **n** | 36 | 36 | 27 | 32 | |
| **Age (mean (SD))** | 65.61 (4.69) | 66.44 (4.75) | 65.22 (4.16) | 65.75 (5.02) | 0.764 |
| **Female sex (%)** | 25 (69.4) | 25 (69.4) | 21 (77.8) | 23 (71.9) | 0.880 |
| **Race (%)** | | | | | 0.798 |
| White | 32 (88.9) | 31 (86.1) | 22 (81.5) | 30 (93.8) | |
| African American/Black | 2 ( 5.6) | 3 ( 8.3) | 3 (11.1) | 2 ( 6.2) | |
| Asian | 2 ( 5.6) | 2 ( 5.6) | 2 ( 7.4) | 0 ( 0.0) | |
| **Adherence (mean % (SD))** | 83 (11) | 81 (12) | 76 (14) | 82 (11) | 0.083 |
| **Education (mean (SD))** | 5.83 (1.23) | 6.19 (1.14) | 5.89 (0.93) | 5.94 (1.22) | 0.564 |
| **Income (mean (SD))** | 7.36 (1.64) | 7.39 (1.68) | 7.22 (1.63) | 8.00 (0.95) | 0.183 |
| **Employment (%)** | | | | | 0.699 |
| FT | 8 (22.2) | 3 ( 8.3) | 6 (22.2) | 9 (28.1) | |
| PT | 5 (13.9) | 6 (16.7) | 1 ( 3.7) | 5 (15.6) | |
| Retired PT | 7 (19.4) | 7 (19.4) | 4 (14.8) | 5 (15.6) | |
| Retired | 15 (41.7) | 19 (52.8) | 15 (55.6) | 12 (37.5) | |
| Unemployed | 0 ( 0.0) | 1 ( 2.8) | 1 ( 3.7) | 1 ( 3.1) | |
| Home maker | 1 ( 2.8) | 0 ( 0.0) | 0 ( 0.0) | 0 ( 0.0) | |

Household income: 9 = Chose not to answer, 8 = >$40,000, 7 =$30,001-$40,000, 6=$25,001-$30,000, 5=$20,001-$25,000, 4=$15,001-$20,000, 3=$10,001-$15,000, 2=$5,001-$10,000, 1= <$5000. PT = part time, FT = full time. Education = number of years.

### 4. Sensitivity analysis

We replicated our pattern of results in just the participants who were randomized to the walking groups only as these are the interventions that are traditionally considered aerobic exercise interventions. Despite a large reduction in the sample size, the pattern of results remains the same. Because the behavioral only model did not produce predictions using elastic net (as in the main paper) we present results for both Ridge regression and Elastic net to allow for more direct comparisons.

Supplementary table 4. Prediction models in the walking and walking + group only

| N=59 | R2 | RMSE | MAE |
|---|---|---|---|
| **Behavioral** | | | |
| Ridge regression | 0.09031898 | 0.1210731 | 0.0992275 |
| Elastic net | N/A | N/A | N/A |
| **Imaging** | | | |
| Ridge regression | 0.1274985 | 0.1185249 | 0.09551711 |
| Elastic net | 0.04349826 | 0.1249851 | 0.09839319 |
| **Multimodal** | | | |
| Ridge regression | 0.239279 | 0.1114668 | 0.09175264 |
| Elastic net | 0.1524032 | 0.1168288 | 0.09512968 |
